# Passive irrigation increases transpiration and reduces heat stress for urban *Lophostemon confertus* trees in metropolitan Sydney

**DOI:** 10.64898/2026.04.29.721794

**Authors:** Davide Siclari, Mark G. Tjoelker, A. Chathurika S. Perera, Sebastian Pfautsch, Paul D. Rymer, Renée M. Marchin

## Abstract

Urban environments typically experience higher air and surface temperatures than surrounding natural landscapes, making urban vegetation crucial for cooling local areas and improving the health of city residents. The long-term survival of young, establishing urban trees may be further constrained in planting contexts where impervious surfaces limit the absorption and retention of precipitation. Here, we investigated how a retrofitted passive irrigation system affected the physiology and growth of young, establishing, *Lophostemon confertus* (brush box) street trees in a hot and dry suburb of western Sydney, Australia. During the 2024-2025 austral summer, three years after planting, there were a total of 16 days above 35 °C, with several periodic dry periods. Irrigated *L. confertus* trees had higher water availability (i.e., higher predawn leaf water potential, *Ψ_pre_*), lower water stress (i.e., higher midday leaf water potential, *Ψ_mid_*, more frequently above turgor loss point), greater stomatal conductance (*g_s_*) on hot and dry summer days, and reduced leaf temperatures (*T_leaf_*), compared to control trees. No significant differences in growth rates were observed between irrigated and control trees during the first three establishment years, but irrigated trees had greater crown survival during the hot, dry summer. Our results suggest passive irrigation may mitigate heat and drought stress in urban trees by increasing water access to support transpiration that prevents leaves from overheating, maintaining tree health. By sustaining cooler and healthier urban tree canopies during extreme summer conditions, passive irrigation may strengthen the climate resilience and liveability of cities under future warming.

## 1 Introduction

The world’s forests and urban environments are increasingly exposed to extreme heatwaves, with temperatures even exceeding 50 °C in some of the hottest cities, threatening both humans (WHO, 2024) and urban trees (Marchin et al., 2022a, 2022b). As a result of the higher cover of impervious surfaces and limited vegetation in cities, surface temperatures in urban areas are 2 to 15 °C warmer than the surrounding countryside (Gómez-Baggethun and Barton, 2013; Duinker et al., 2015; Cueva et al., 2022; McPhearson et al., 2023, 2025). Globally, urban vegetation is one of the most widespread solutions to decrease temperatures and improve human thermal comfort (Saaroni et al., 2018; Wai et al., 2022). Through transpiration, urban vegetation can reduce air temperatures by 1-3 °C at the whole city scale (Livesley et al., 2016; Sanusi et al., 2020). Trees also effectively reduce the surface temperatures of urban streets and footpaths, which can reach 84.5 °C in unshaded areas, whereas shaded surfaces are 7 to 45 °C cooler (Napoli et al., 2016; Sharmin et al., 2023a, b). Heatwave conditions have recently caused extensive canopy dieback and mortality of urban vegetation (Marchin et al., 2022a; Esperon-Rodriguez et al., 2025a), with greater vulnerability of short-lived deciduous species (Zhang et al., 2025) and of young trees (Marchin et al., 2025; Lalor et al., 2023; Lv, Gangwisch and Saha 2024; Jääskeläinen et al., 2025; Parhizgar et al., 2025).

Urban street trees might be predisposed to drought stress as a result of urban infrastructure and planning, including high percentages of impervious surfaces (Marchin et al., 2025), compacted soils (Asif et al., 2023), and restricted rooting volumes (Sand et al., 2018; Zhang et al., 2025). Impervious surfaces exacerbate surface runoff, reducing water availability for urban vegetation and enhancing transport and accumulation of dissolved pollutants (Jacobson, 2011). Compacted soils with poor infiltration promote rapid flooding during large rainfall events (Morgenroth, 2013), heightening the risk of root rot in newly established trees (Day et al., 2010), while the lack of irrigation in dry periods risks exposing urban trees to limited water availability. High percentages of xylem cavitation (Savi et al., 2025) and drought-induced growth reductions (Nitschke et al., 2017; Marchin et al., 2025) have been reported for trees that are surrounded largely by impervious surfaces. The frequency, severity, and length of water limitation are dependent on local climate: in mesic cities, trees rely primarily on rainfall for their dominant water supply (Smith et al., 2024), whereas trees in arid and semi-arid environments depend largely on irrigation (Bijoor et al., 2012; Gómez-Navarro et al., 2019). Water access is critical for the tolerance of heatwaves (Taiz et al., 1991; Meineke et al., 2018; Marchin et al., 2022b), as the process of transpiration effectively cools tree leaves. Newly established and small trees are particularly vulnerable to prolonged dry periods in urban environments (Haase et al., 2022, Marchin et al., 2025).

Irrigation of urban trees could help protect tree leaves from overheating on hot summer days, plus accelerate tree growth and urban canopy expansion (Montague & Bates, 2015), but it is costly (Pataki et al., 2011) and unfeasible at a large scale for many cities (Luketich, Papuga & Crimmins 2018). The demand for irrigation to sustain urban vegetation is likely to increase by 2050 due to changes in seasonal weather patterns that alter precipitation patterns, including more intense and prolonged droughts (Esperon-Rodriguez et al., 2025). Traditional urban water management practices prioritise the rapid removal of stormwater with minimal infiltration or water retention, which is likely to exacerbate the effects of climate change (EPA, 2013; Climate Council of Australia Ltd., 2018; Thom et al., 2022). Some cities are starting to implement a different approach by using water-storing passive irrigation (PI) systems that capture and redistribute rainfall. These systems require a higher upfront infrastructural cost (CRC, 2020), compared to traditional irrigation methods, but are an innovative solution that could simultaneously reduce the amount of water that needs to be processed by existing stormwater control measures (Morgenroth et al., 2013; Scharenbroch et al., 2016) and divert water to urban vegetation. Out of 109 cities across 21 countries, only 7% currently use PI systems that retain stormwater for green infrastructure (Esperon-Rodriguez et al., 2025).

Previous studies have documented that the effects of passive irrigation on tree growth can vary with PI design (EPA, 2013), tree age (Thom et al., 2022; Ow and Chan, 2021; Szota et al., 2019), and tree species (Grabosky and Bassuk, 2016; Gleeson et al., 2022). Irrigated trees can have up to 2- to 6-fold higher growth than control trees during their establishment (Grey et al., 2018; Ow and Chan, 2021), but growth of established trees may not benefit from passive irrigation (Szota et al., 2019). Fast-growing species, such as *Melia azedarach* (Chinaberry tree) and *Acer campestre* (field maple), had 44-65% greater growth with passive irrigation (Gleeson et al., 2022; Thom et al., 2022). It is necessary that PI systems avoid waterlogged conditions to promote tree growth, although this increases the potential for drought stress when rainfall is scarce (Grey et al., 2018; Ow and Chan, 2021). Potentially, PI systems can reduce drought stress of urban trees, relative to unirrigated trees, by increasing local soil moisture (Armson et al., 2013; Rahman et al., 2017). Greater soil moisture supports higher stomatal conductance and photosynthetic rates (Gobatti et al., 2025), which can theoretically increase canopy cooling and urban tree growth by preventing stress and mortality during extreme heat events. There is a need for detailed physiological studies that evaluate the benefit of PI systems for tree function and survival to justify their greater infrastructural cost and to support wider deployment of these innovative systems.

In many Australian cities, precipitation is highly variable and fluctuates between periods of mild to intense drought, along with episodic high-intensity storms, so managing water sustainably is particularly pressing. This study utilised a PI system that had been retrofitted into a residential street in western Sydney, one of the hottest and most rapidly developing urban areas in Australia. Many suburbs in this region are characterized by low canopy cover (<20%) and a pronounced urban heat island effect of approximately 6-12 °C (Pfautsch et al., 2024; NSW Government, 2025). The aim of our study at this demonstration site was to determine if and how the PI system affected the physiology, growth, and survival of the most widely planted native evergreen tree species in Australian cities (brush box, *Lophostemon confertus*; Frank et al., 2006; Esperon-Rodriguez et al., 2021). In the third year after planting, the austral summer of 2024-2025, the local climate was periodically hot and dry, with thirteen days above 35 °C, three days exceeding 40 °C, and a prolonged dry period of 26 days with negligible rainfall (≤15 mm) at the peak of summer. In this study, we compared irrigated street trees to both unirrigated control trees and nearby park trees, addressing the following research questions:

- Do PI systems increase the health and growth of urban trees during the early establishment years (e.g., in the first three years after planting)?
- Does additional water captured by PI systems mitigate heat and drought stress in urban trees?
- Do PI systems decrease leaf temperatures in urban trees?

We hypothesised that, relative to control trees and nearby park trees, irrigated trees will have greater growth, higher water availability (e.g., higher predawn leaf water potential, *Ψ_pre_*), less midday water stress (e.g., higher midday leaf water potential, *Ψ_mid_*), greater stomatal conductance (*g_s_*), and cooler leaf temperatures (*T_leaf_*).

## 2 Materials and methods

### 2.1 Study site

The study took place on an urban residential street in western Sydney - Wilkes Crescent (Tregear) within the Blacktown local government area of Sydney, New South Wales, Australia (Fig. S1). Blacktown has a population of 460,000 residents, situated approximately 34 kilometres west of central Sydney at an elevation of 78 meters above sea level. Blacktown has a subtropical climate with a mean annual temperature of 16.9 °C and a mean annual precipitation of 706 mm. The hottest and coldest months are January (average daily maximum 29.8 °C ± 2.4, minimum 18 °C ±1.9 °C) and July (average daily maximum 17.4 °C ± 0.9, minimum 5.9 °C ± 0.9), respectively. The wettest and driest months are February (121.6 mm ± 103.6) and September (38.3 mm ± 29.5), respectively. Long-term climate data was averaged from 1996-2026 from the Australian Bureau of Meteorology station 067119 at Horsley Park, located 10 km from the study site.

Environmental conditions were monitored on site via the ATMOS 41W weather station (METER Group, UT, USA) installed at three meters from the ground to measure precipitation, air temperature (min, max, average), relative humidity, wind speed and direction, and maximum wind gust every 15 minutes during the 2024-2025 austral summer. Soil moisture was measured by six 90-cm sensors, including two control (N = 2 sensors, as one sensor was excluded due to technical issues) and three irrigated (N = 3 sensors, Sentek Drill & Drop, Cheltenham, VIC). Soil moisture data from a depth of 45 cm was used for analysis. Relative extractable soil water content (REW) was calculated using the following equation (Eq. 1):

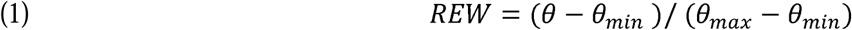

Where *θ* is the soil water content, *θ_min_* is the minimum soil water content and *θ_max_* is the mean maximum soil water content over the 2024-2025 austral summer. Readings from three peak rainfall events were averaged to determine *θ_max_*; each reading was taken two hours after the rainfall event to account for soil macropore drainage and avoid overestimation of *θ_max_* Soil moisture data was unavailable after 16 January 2025, when a storm with >80 mm of rain fell within two hours, and the underground sensor box suffered water infiltration.

### 2.2 Passive irrigation system

The passive irrigation (PI) system was retrofitted into a street bordered by residential housing adjacent to a recreation area (Fig. S1c, e). It was created as a trial demonstration site before the larger deployment of similar PI systems testing four tree species on six urban streets; however, these other retrofitted sites were not constructed due to post-COVID cost escalation. This retrofitted PI system captured stormwater through four street-level inlets from a catchment area of 426.56 m^2^, stored it in a gravel trench, and was connected to the existing stormwater management system to allow discharge of any overflow and prevent waterlogging (Fig. S1d). Four street-level inlets directed the water underground into a polymer stormwater sediment collection pit. The water was then conveyed into a perforated 100-mm PVC pipe positioned at the bottom of a trench that was 95 m long, 45 cm in width and 65 cm in depth. The trench held 9756 L of water at full capacity due to several layers of geofabric textile material, rocks, and metal nets. Vertical PVC pipes were installed for monitoring equipment. The first and last trees were within 1.5 m from the catching and discharging pits, spaced 101.8 m apart.

Twenty-five trees of *L. confertus* (brush box) were planted at approximately six-metre spacing along the street in November 2021. *Lophostemon confertus* is native to eastern Australia, naturally distributed from northeastern New South Wales to southeastern Queensland, where it typically occurs in coastal and subcoastal rainforests, and is commonly planted throughout Australia’s major cities. There were twenty-two irrigated trees with access to the stored stormwater, which were staggered on either side of the underground trench, while the last three trees on the street served as controls without access to stored stormwater. Control trees were necessarily spatially separated from irrigated trees to ensure their tree roots could not access the stored stormwater in the underground trench. Tree age at time of planting was 16-18 months in 45-L bags (Table S2). Site maintenance included regular cleaning of the inlets, grass mowing and basal pruning. Trees were manually irrigated in their establishment year, but irrigation ceased in February 2022. Another eleven control trees of the same species were located within a nearby open grassy area (Shanes Park Reserve, 4 km away; Fig. S1f) and did not have access to any passive irrigation system. Tree spacing, age at planting, and maintenance history were similar to the Wilkes Crescent trees, as all trees were 12-16 months at time of planting and were planted in September 2020.

### 2.3 Tree growth and crown dieback

Growth measurements across all three tree groups (N = 22 irrigated, N = 3 control, N = 11 park) included height, crown width and length, and stem diameter both at the base of the tree and at breast height (DBH, 1.3m). Growth was measured in February and November 2022, September 2023, December 2024 and March 2026. Crown dieback was visually assessed in September 2023 and December 2024, considering the difference between the percentage of missing, scorched and/or desiccated leaves, relative to the natural crown shape of the species. Crown size was estimated as ground-projected crown area measuring two perpendicular diameters at the base of the crown and calculated using the equation for the area of an ellipse (Eq. 2). Relative growth rates (RGR, Eq. 3) and crown survival (Eq. 4) were calculated using the following equations:

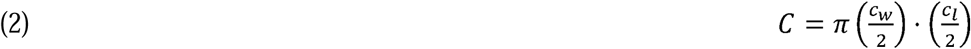

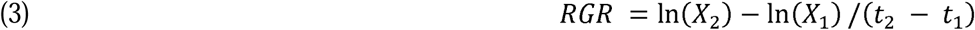

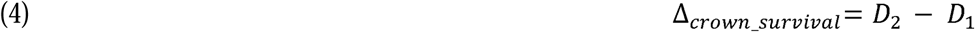

where C is total canopy area, c_w_ and c_l_ canopy width and length, respectively. X_1_ is the initial height, canopy area or diameter of the tree at time t_1_, and X_2_ is the final height, canopy area or diameter of the tree at time t_2_. RGR were calculated for all growth parameters. Crown survival was calculated as the difference between the crown dieback in 2024 (D_2_) minus the dieback in 2023 (D_1_).

Root structure and rooting depth were assessed on 18 March 2025 in 50-cm soil cores removed from the base of the first, fourth and eighth irrigated trees. A second excavation of equal depth was done to intercept the trajectory of detected roots extending toward the trench. Irrigated trees had many shallow roots near the soil surface that extended into the side walls of the water-storing trench (Fig. S2a, b).

### 2.4 Leaf morphology

We measured plant functional traits related to drought or heat tolerance, including leaf size, leaf thickness, leaf mass area (LMA, Eq. 5), and leaf dry matter content (LDMC, Eq. 6). Leaf traits (N = 3 leaves per tree) were measured on fully-expanded leaves for both irrigated (N = 8) and street control trees (N = 3) in March 2025. Leaf size was measured using a flatbed scanner and ImageJ software (Schneider et al., 2012). Thickness was measured at the top, middle and bottom of the leaf blade, avoiding the midrib and veins, with digital callipers. Leaves were rehydrated for 1 hour using the standing rehydration technique (Arndt et al., 2015), weighed to obtain fresh weight, then oven-dried for 48 h at 70 °C to a constant weight to obtain dry mass.

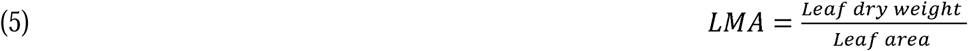

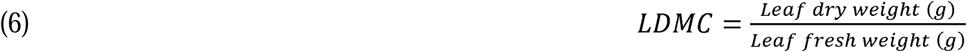

### 2.5 Tree water relations

All three groups (i.e., irrigated, control, park) of *L. confertus* were measured monthly during the moderately wet autumn 2024 (March to June 2024; Table S1). Thereafter, only trees within the PI system were monitored during the austral summer of 2024-2025 (November 2024 to February 2025). On nine sunny measurement days (roughly 3 days per summer month, DJF; Fig. S3), we measured midday leaf water potential (*Ψ_mid_*) by collecting leaves between 13:00 and 14:00. Predawn leaf water potential (*Ψ_pre_*) was measured one day per summer month by collecting leaves between 4:00 and 5:00. Measurements were done on two sun-exposed and fully mature leaves per plant on the first eight irrigated trees at the beginning of the line, furthest from the three controls positioned at the end of the system. Excised leaves were placed in an iced portable cooler inside sealed and humidified (via wet paper towel) plastic bags using the double-bag method (Rodriguez-Dominguez et al., 2022). Once in the laboratory, leaves were transferred into a refrigerator at 3.4 °C and kept cool and dark until equilibrated (1 hour). We measured *Ψ* using a pressure chamber (Model 1505D, PMS Instruments, Corvallis, OR, USA). *Ψ* values were determined as the mid-point between the first sign of water emerging from the petiole, and the flooding of the entire xylem area, using a dissecting microscope with fibre-optic lights to clearly observe the balance pressure at the cut end of the petiole. All leaves were processed within 2-3 hours from collection.

To estimate the leaf water potential at turgor loss point (TLP; Bartlett et al. 2012), leaves (N = 1 leaf per tree) on the same irrigated (N = 8) and control trees (N = 3) were collected at 14:00 on 23-27 February 2025 with an analogous method to the *Ψ* measurements. Leaves were rehydrated for 1-2 hours via the standing rehydration technique (Arndt et al., 2015), considering fully rehydrated leaves to have *Ψ* ≥ − 0.3 MPa. Leaves were cut into disks avoiding the midrib, wrapped in foil, and submerged in LN_2_ for 2 minutes, then re-equilibrated in a humidified plastic bag for ten minutes. Leaves were pierced with forceps before measuring osmotic potential (π*_o_*) using an osmometer (WP4C, Decagon Devices Inc., Pullman, WA, USA) until equilibrium was reached (<0.01 MPa change in osmotic potential over 2 min). TLP was calculated using the equation from Bartlett et al. (2012) (Eq. 7):

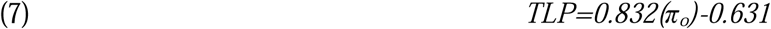

### 2.6 Leaf thermoregulation and thermal tolerance

Leaf temperature (*T_leaf_*), quantum yield of photosystem II (□_PSII_) and stomatal conductance (*g_s_*) were measured during sunny days, including one extreme (maximum *T_air_* ≥ 40 °C), two hot (maximum *T_air_*= 35-40 °C) and six average days (maximum *T_air_* = 30-35 °C) using a porometer/fluorometer (LI-600, LICOR Biosciences, Lincoln, NE, USA). Measurements were made on four sun-exposed and fully mature leaves per tree on the same irrigated (N = 8) and control trees (N = 3). The park trees (N = 11) were also measured monthly during autumn 2024. *T_leaf_* was measured at the hottest time of day (13:00-15:00), as determined by preliminary testing. To measure _PSII_, we measured steady-state fluorescence (*F*_s_) and then applied a saturation pulse to measure maximum fluorescence in the light (*F*_m_′). Transpiration (*J*) was calculated using the equation (Eq. 8):

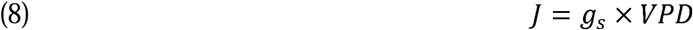

where *g_s_* is stomatal conductance, and VPD is vapor pressure deficit, defined as the difference between the amount of moisture in the air and the amount it can hold at full saturation.

We determined tree canopy temperature for all 25 trees using a thermal infrared camera (Model T640, FLIR, Wilsonville, OR) on 13 March 2025 (maximum *T_air_* = 41.1 °C) to better understand spatial patterns in temperature across the tree canopy. Images were captured between 13:00 and 15:00 from the north-facing side of the tree. In addition, we quantified surface temperature for different types of ground cover (asphalt, sunlit versus tree-shaded areas, bare grass, car surfaces, etc.).

Leaf critical temperature (*T_crit_*) was determined by measuring the temperature-dependent rise of steady-state chlorophyll α fluorescence (T-F_0_) on one fully mature and sun-exposed leaf (N = 1 leaf per plant; N = 8 irrigated, N = 3 control trees; Schreiber and Berry, 1977). Leaves were collected at midday between 1 and 7 February 2025 and kept in the dark. Immediately after detachment, samples were placed in humidified plastic bags containing a moistened paper towel, sealed, and transported to the laboratory in a chilled portable cooler. Upon arrival, leaves were stored at 3.4 °C and processed within 24 hours of collection. Leaves were pressed firmly on top of a Peltier thermoelectric cooler (Model APH-161-12-18-E, European Thermodynamics, Kibworth, UK) inside a custom-built leaf cuvette. A Peltier temperature controller (Arduino Nano V3, Monza, Italy) regulated temperature changes via a PID control algorithm and was connected to a touch-screen computer interface (Model 3B, Raspberry Pi, UK). Two identical thermistors monitored and recorded the temperatures of both the abaxial and adaxial leaf surfaces (Model MC65F103A, GE Sensing/Thermometrics, Billerica, MA, USA). These two temperatures were then averaged to calculate *T_leaf_*. In the clamping device, at a 60° angle to the leaf surface and through a fiber-optic cable, the leaves were exposed to a low-intensity, far-red illumination (<1 μmol m^-2^ s^-1^) to maintain PSII in an oxidised state (Valladares and Pearcy, 1997). Steady-state leaf fluorescence was recorded every 1 second via a fluorometer (MINI-PAM-II/B, Heinz Walz GmbH) while the Peltier temperature increased by 1 °C every minute from 30 °C. Measurements were continued until the F_0_ began to decline after reaching its maximum value, or the Peltier temperature reached 65 °C. *T_crit_* was then calculated as the intersection of the linear slow- and fast-rise phases of the T-F_0_ curve (Knight and Ackerly, 2002; Schreiber and Berry, 1977). The slow-rise phase was calculated using F_0_ from 30° to 40 °C, while the fast-rise phase using F_0_ from ± 1.5 min of the mid-point between the minimum and maximum F_0_ values. Thermal safety margins (*TSM*) were calculated as the difference between the *T_crit_* and the maximum recorded leaf temperature on 28 January 2025 (maximum *T_air_*= 40.2 °C), within a week of the *T_crit_* measurements.

### 2.7 Statistical analysis

To determine if *T_leaf_*, *Ψ*, *g_s_*, □_PSII_, and soil moisture differed between irrigated, control, and park trees, linear mixed-effects models were used, with date and tree identity included as random factors to account for repeated measures. Relative extractable water content (REW) differences between treatments were evaluated using analysis of variance (ANOVA). An irrigation effect on the relationships between hydraulic safety margin and loss of function (TLP *-Ψ_mid_*) and rehydration capacity and water access (*Ψ_pre_* - *Ψ_mid_*), were tested using soil VWC as a covariate. A logarithmic regression was used to describe the relationship between soil VWC and Ψ. RGR were tested among treatments, dates and interaction using ANOVA. Linear and nonlinear regressions were tested to determine the relationships between quantum yield of photosystem II (□_PSII_) and *T_leaf_*, *g_s_* and VPD, and *T_leaf_*and *T_air_*; Akaike’s information criterion was used to select best fitting models. Tukey’s Honestly Significant Difference (HSD) test and Wilcoxon tests were applied to identify differences between irrigated and control plants. All data were tested for normality using the Shapiro-Wilk test. Thermal safety margins, *T_leaf_* and *Ψ_mid_* were log-transformed to achieve normality. Means were considered significantly different at *p* ≤ 0.05; errors were expressed as SE. Statistical analyses were computed using R studio version 4.4.3 (R Core Team, 2025).

## 3 Results

### 3.1 Environmental conditions

The 2024-2025 austral summer in western Sydney was periodically hot and dry, although there was a two-week wet period in mid-January with high rainfall (Table 1, Fig. S3). The peak of summer was on 7 January 2025 (maximum *T_air_* = 38 °C, maximum *T_leaf=_* 47.48 °C), following a twenty-six-day period with little to no rainfall (<15 mm). The maximum air temperature was 42.2 °C, recorded on 29 January 2025 along with a maximum VPD of 8.3 kPa. Passive irrigation increased daily mean REW by 13% over the period from December 2024 to mid-January 2025 for irrigated trees (F_1,55_ = 12.87, *p* <0.001; Fig. S3b).

**Table 1.** Monthly and 30-year climate statistics. Total monthly precipitation and mean maximum monthly air temperature (*T_max_mean_*) for the 2024-2025 austral summer (DJF), plus the warm months of November and March, relative to 30-year (1996-2026) mean precipitation in the western Sydney suburb of Tregear, NSW, Australia. Monthly data for the 2024-2025 summer were available from a weather station on site, while the 30-year climate means are from a Bureau of Meteorology site (BoM Horsley Park station, 067119) located 10 km from the study site.

| Month | 2024-2025 precipitation<br>(mm) | 30-Year<br>precipitation (mm) | 2024-2025<br>$T_{max\_mean}$ (°C) | 30-Year $T_{max\_mean}$<br>(°C) |
| --- | --- | --- | --- | --- |
| November | 63.4 | 75.5 | 27.8 | 26.5 |
| December | 63.3 | 65.6 | 31.4 | 28.5 |
| January | 168.7 | 79.3 | 29.6 | 29.8 |
| February | 91.8 | 121.6 | 29.8 | 28.7 |
| March | 86.8 | 89.6 | 27.6 | 26.8 |

### 3.2 Tree growth and survival

There was no significant difference between irrigated and control trees in RGR for tree height (F_1,23_ = 0.811, *p* = 0.377), DBH (F_1,23_ = 2.31, *p* = 0.14), or basal growth (F_1,23_ = 1.30, *p* = 0.266) for any study year (Table 2). There was significantly higher canopy area RGR for irrigated trees (F_1,23_ = 4.39, *p* = 0.047), relative to control trees, in the second year (2022-2023) but not in the third or fourth years (2023-2024, 2024-2026; Table 2). Trees had greater crown dieback in 2024 than in 2023 (F_1,23_ = 8.12, *p* < 0.01), with irrigated trees maintaining significantly lower crown dieback compared to controls (F_1,23_ = 6.04, *p* = 0.021; Fig. 1). Irrigated trees had bigger leaves than controls (F_1,9_ = 16.26, *p* < 0.01), but leaves did not differ in thickness (F_1,9_ = 0.76, *p* = 0.4), LMA (F_1,9_ = 0.44, *p* = 0.51), or LDMC (F_1,9_ = 2.91, *p* = 0.12; Fig. 2).

**Figure 1.**
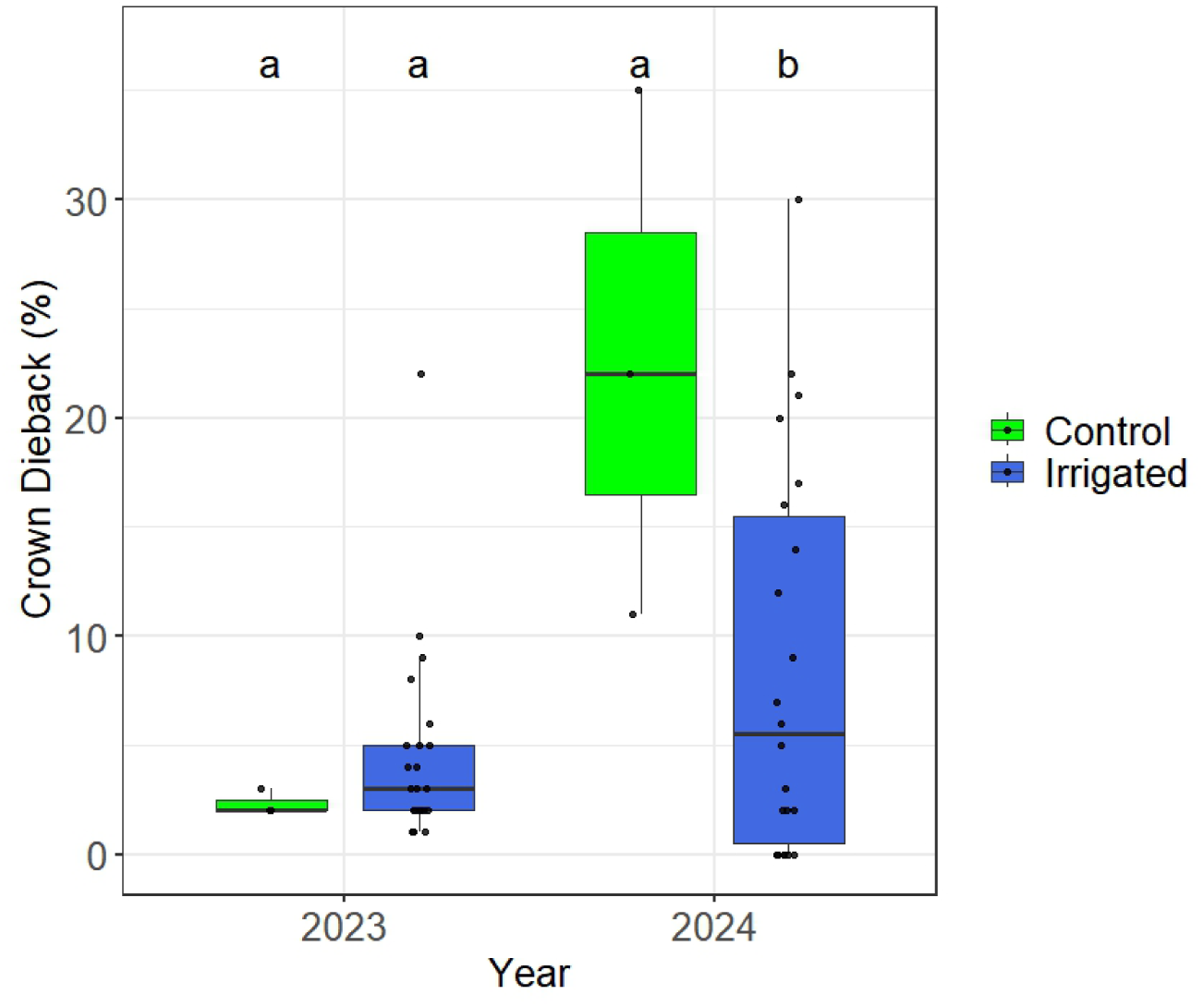
The change in crown dieback between September 2023 and December 2024 for irrigated (N = 22) versus control (N = 3) trees of *Lophostemon confertus* in the western Sydney suburb of Tregear, NSW, Australia. Irrigated trees are coloured blue, controls in green. Means not connected by the same letter are significantly different (Tukey honestly significant difference, *p* < 0.05). Error bars represent ± standard error of the mean; whiskers indicate 1.5 × interquartile range. Points indicate tree level data.

**Figure 2.**
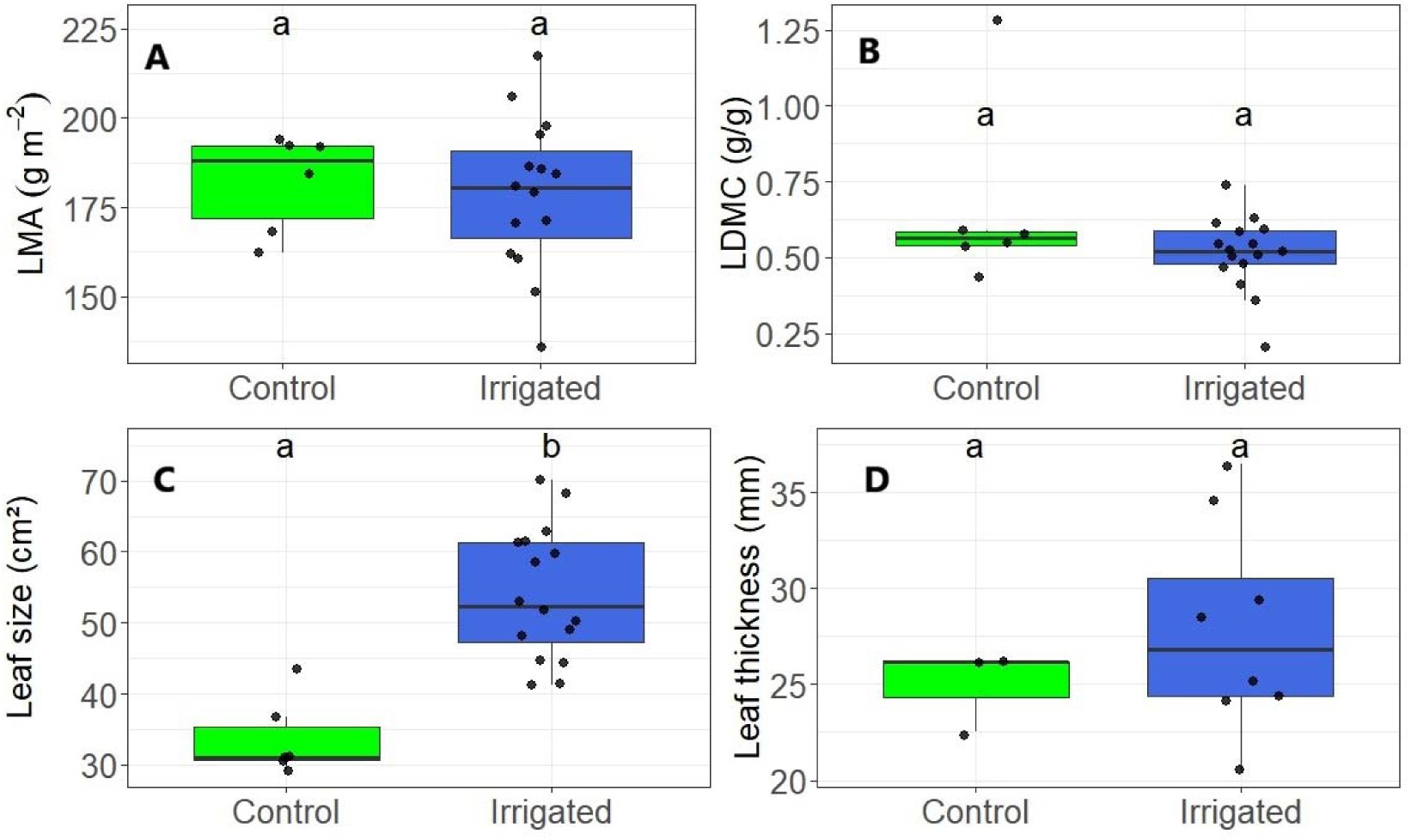
Comparison of leaf (N = 2 leaves per tree) morphological traits among irrigated (N = 8) and control (N = 3) trees of *Lophostemon confertus* in the western Sydney suburb of Tregear, NSW, Australia, including (A) leaf mass per area (LMA), (B) leaf dry matter content (LDMC), (C) leaf size and (D) thickness. Irrigated trees are coloured blue, controls in green. Means not connected by the same letter are significantly different (Tukey honestly significant difference, *p* < 0.05). Error bars represent ± standard error of the mean; whiskers indicate 1.5 × interquartile range. Points indicate tree level data, and the average between two leaves in panel D.

**Table 2.** Tree growth. Mean relative growth rates for irrigated (N = 22) and control (N = 3) trees of *Lophostemon confertus* for height, diameter at breast height (DBH, 1.3m), basal diameter (trunk diameter at 10 cm from the ground) and canopy area (m^2^) since establishment in the western Sydney suburb of Tregear, NSW, Australia in 2021. All means are expressed in cm cm^-1^ y^-1^ (± SE). Yearly means not connected by the same letter are significantly different (Tukey honestly significant difference, *p* < 0.05). Shading groups the measurements relative to the same year. Year since planting (YSP) is also provided.

| Year | YSP | Measurement | Irrigated (cm cm<sup>-1</sup> y<sup>-1</sup> ± SE) |  |  | Control (cm cm<sup>-1</sup> y<sup>-1</sup> ± SE) |  |  |
| --- | --- | --- | --- | --- | --- | --- | --- | --- |
| 2022-2023 | 2 | RGR_Height | 0.145 | ±0.067 | a | 0.157 | ±0.066 | a |
| 2022-2023 | 2 | RGR_DBH | 0.003 | ±0.002 | a | 0.004 | ±0.0006 | a |
| 2022-2023 | 2 | RGR_Basal | 0.005 | ±0.002 | a | 0.006 | ±0.001 | a |
| 2022-2023 | 2 | RGR_Canopy_Area | 0.392 | ±0.07 | a | 0.170 | ±0.054 | b |
| 2023-2024 | 3 | RGR_Height | 0.180 | ±0.079 | a | 0.207 | ±0.072 | a |
| 2023-2024 | 3 | RGR_DBH | 0.007 | ±0.001 | a | 0.006 | ±0.001 | a |
| 2023-2024 | 3 | RGR_Basal | 0.008 | ±0.001 | a | 0.007 | ±0.0005 | a |
| 2023-2024 | 3 | RGR_Canopy_Area | 0.386 | ±0.05 | a | 0.531 | ±0.055 | a |
| 2024-2026 | 4 | RGR_Height | 0.170 | ±0.061 | a | 0.174 | ±0.062 | a |
| 2024-2026 | 4 | RGR_DBH | 0.005 | ±0.0006 | a | 0.005 | ±0.0002 | a |
| 2024-2026 | 4 | RGR_Basal | 0.007 | ±0.0009 | a | 0.006 | ±0.0007 | a |
| 2024-2026 | 4 | RGR_Canopy_Area | 0.166 | ±0.03 | a | 0.046 | ±0.05 | a |

### 3.3 Tree water relations

Irrigated trees consistently had higher predawn (*Ψ_pre_*) and midday (*Ψ_mid_*) leaf water potentials than control trees throughout autumn 2024 (F_2,17_ = 7.11, *p* < 0.01; Fig. 3) and the 2024-2025 austral summer (F_1,29_ = 12.58, *p* = 0.001; F_1,9_ = 7.81, *p* = 0.02, respectively; Fig. 4). Both the *Ψ_pre_* and *Ψ_mid_*of irrigated trees were equivalent to nearby park trees in autumn 2024 (Fig. 3). The *Ψ_mid_* of irrigated and park trees were more frequently above their turgor loss point (TLP = −2.21 MPa), while control street trees were often at or below their TLP (Fig. 3, 4b). Stomatal conductance (*g_s_*) did not differ consistently between irrigated and control trees when averaged across the whole 2024-2025 austral summer (F_1,9_ = 3.12, *p* = 0.11, Fig. 5a, S4a), but irrigated trees had higher *g_s_* than park trees during autumn 2024 (F_2,16_ = 5.36, *p* < 0.01; Fig. 3c) and control trees on three hot and dry summer days (REW< 0.1, F_1,9_ = 28.14, *p* < 0.001, Fig. 5b). When hot days (maximum *T_air_* > 30 °C) occurred after recent large rainfall events resulting in wetter soil (REW > 0.3), there was no difference in *g_s_* between irrigated and control trees (F_1,9_ = 0.76, *p* = 0.56, Fig. 5c). Stomatal conductance of all trees decreased with increasing VPD (R^2^ = 0.43, *p* < 0.001), as expected, but *g_s_*-VPD relationships did not differ between treatments (F_1,151_ = 0.38, *p* = 0.54; Fig. S4b). Stomatal conductance was not related to midday leaf water potential across all trees (F_2,7_ = 2.4, *p* = 0.16), but three irrigated trees (tree n. 3, 4 and 6) exhibited consistently higher *g_s_*rates than the other study trees across 2024-2025 austral summer (Fig. S4c). Transpiration was not related to air temperature for both irrigated and control trees over the whole 2024-2025 austral summer (F_3,14_ = 0.68, *p* = 0.57, data not shown). Irrigated trees had high transpiration rates on hot, dry days (REW< 0.1, Fig. S5a), while all trees displayed stomatal closure on milder summer days after recent rainfall (Fig. S5b).

**Figure 3.**
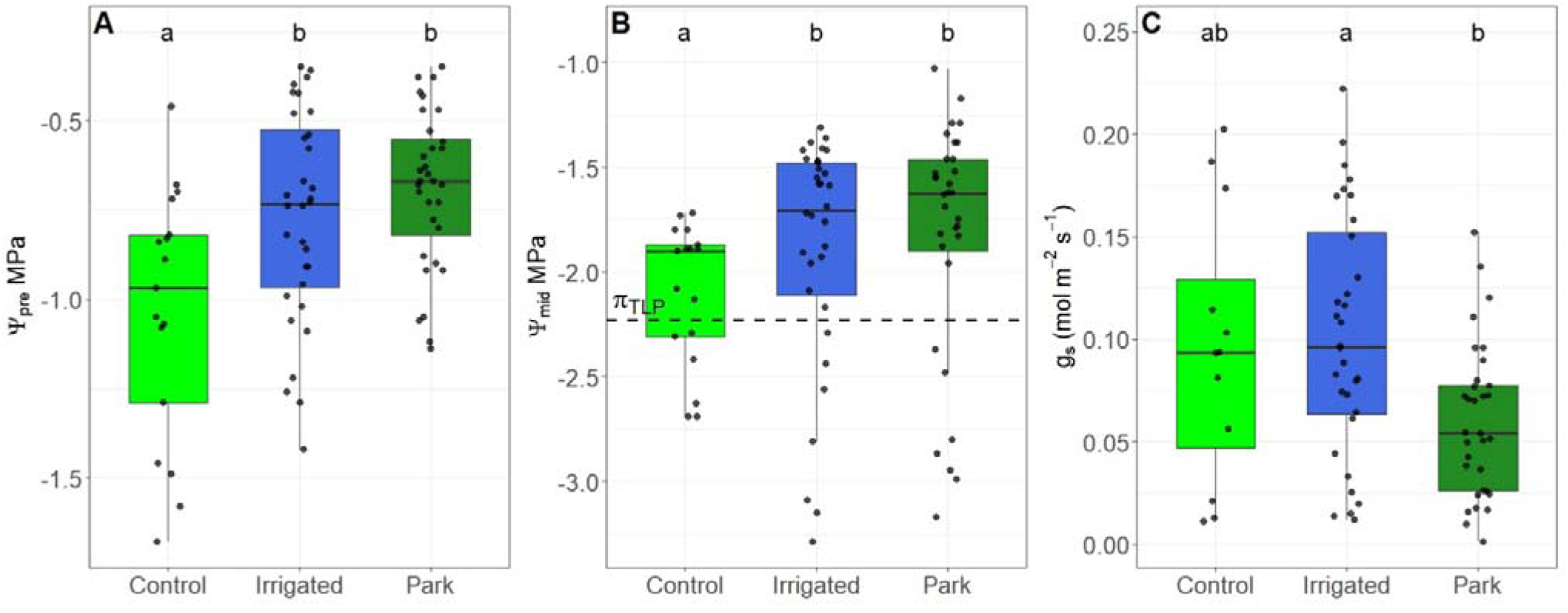
(A) Predawn (*Ψ_pre_*), (B) midday leaf water potentials (*Ψ_mid_*), and (C) stomatal conductance of irrigated (N = 8), control (N = 3), and park trees (N = 11) of *Lophostemon confertus* in Blacktown, during the wet austral autumn 2024. Means not connected by the same letter are significantly different (Tukey honestly significant difference, *p* < 0.05). Error bars represent ± standard error of the mean; whiskers indicate 1.5 × interquartile range. Points are tree-level means (n=4 leaves per tree) from four sampling dates. The horizontal dashed line in panel B indicates the species critical hydraulic threshold or leaf turgor loss point (TLP).

**Figure 4.**
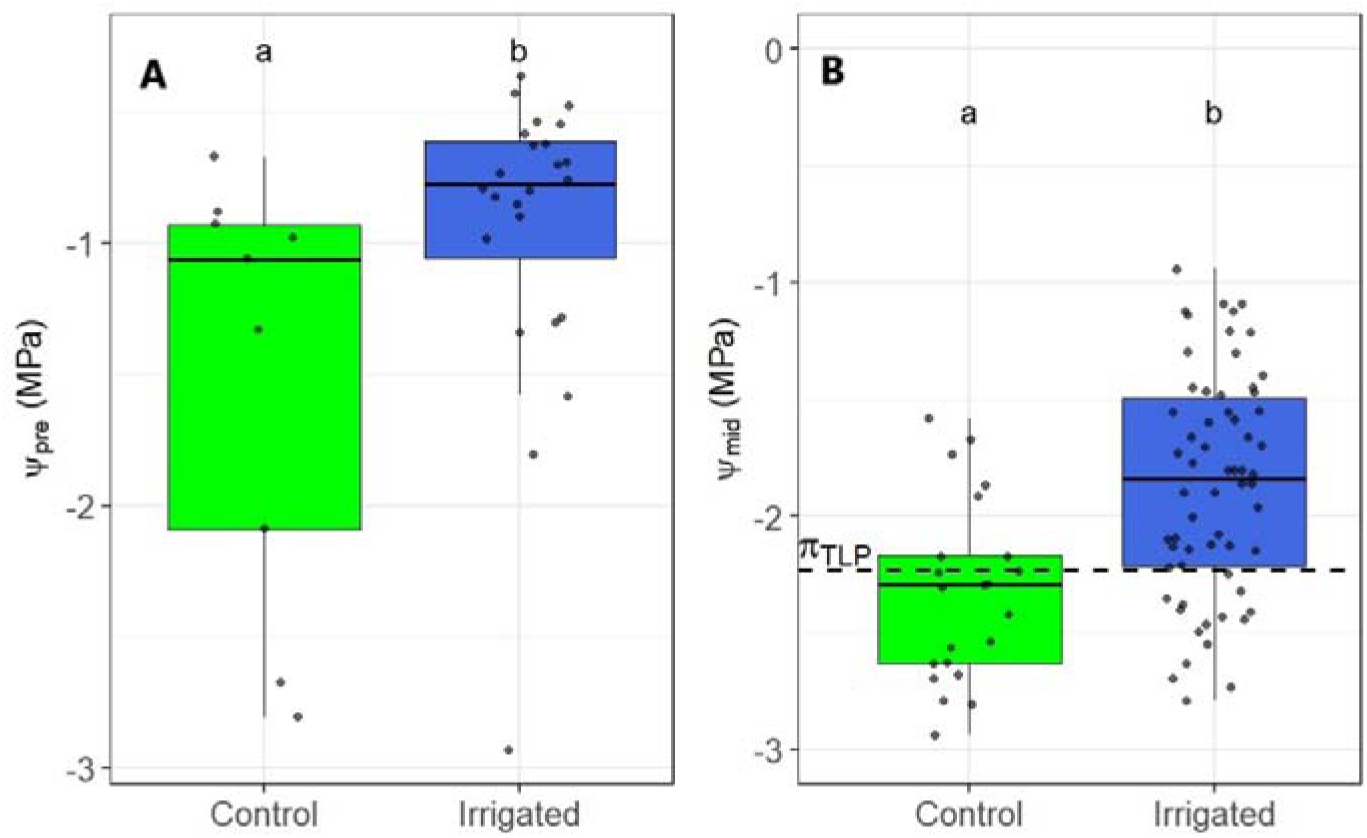
(A) Predawn (*Ψ_pre_*) and (B) midday leaf water potentials (*Ψ_mid_*) of irrigated (N = 8) and control (N = 3) trees of *Lophostemon confertus* in Tregear during the austral 2024-2025 summer. The horizontal dashed line indicates the species critical hydraulic threshold or leaf turgor loss point (TLP). Irrigated trees are coloured blue, controls in green. Means not connected by the same letter are significantly different (Tukey honestly significant difference, *p* < 0.05). Error bars represent ± standard error of the mean; whiskers indicate 1.5 × interquartile range. Points indicate raw data.

**Figure 5.**
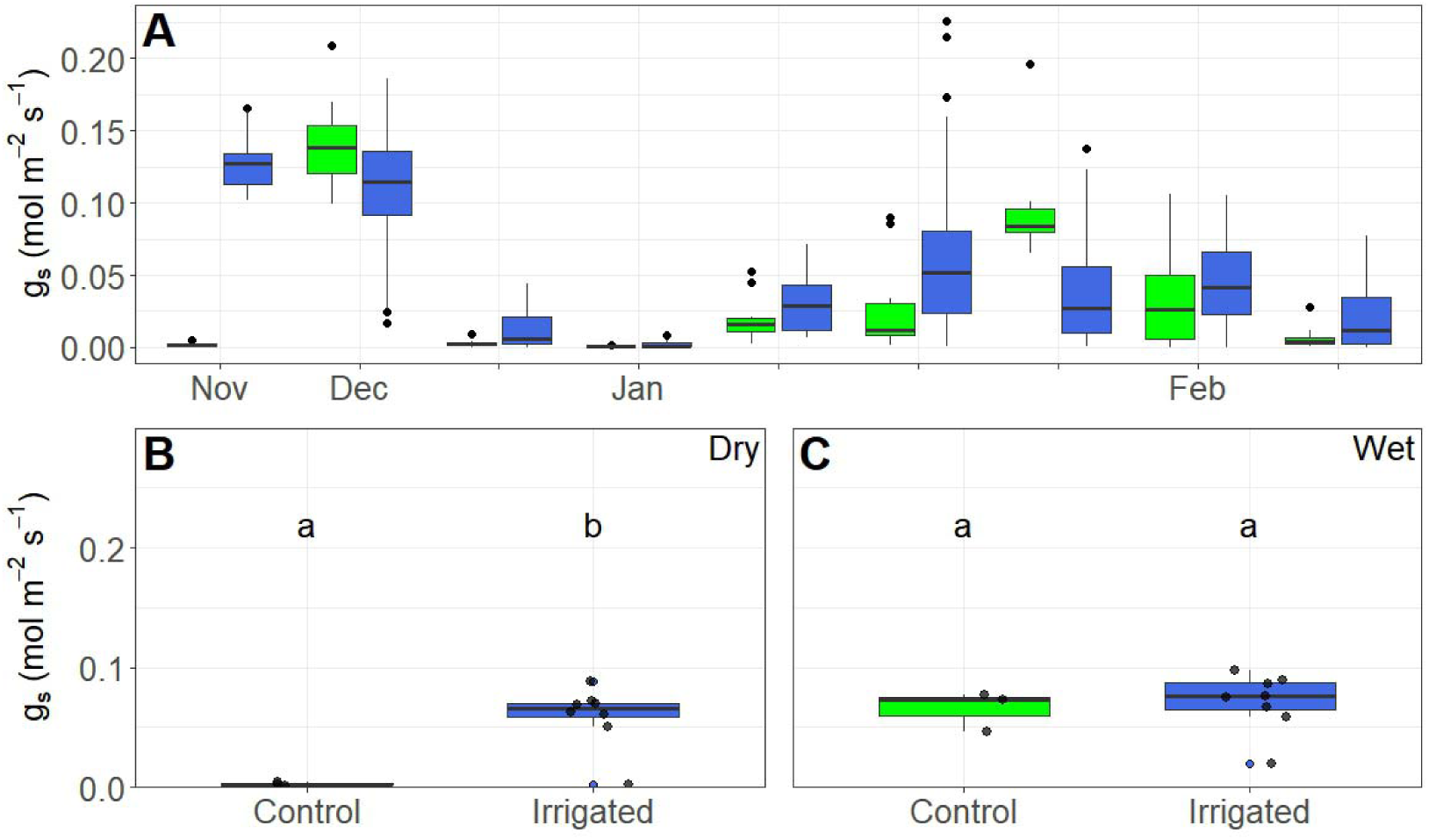
(A) Comparison of stomatal conductance between irrigated (N = 8) and control (N = 3) trees of *Lophostemon confertus* in the western Sydney suburb of Tregear, NSW, Australia during the austral 2024-2025 summer. Points indicate maximum and minimum values. Subset of (B) three dry days when REW <0.1 and (C) three wet days when REW was >0.3. Each point is one individual tree (N = 4 leaves per tree). Irrigated trees are coloured blue, controls in green. Means not connected by the same letter are significantly different (Tukey honestly significant difference, *p* < 0.05). Error bars represent ± standard error of the mean; whiskers indicate 1.5 × interquartile range.

### 3.4 Leaf temperature

Irrigated trees had significantly lower mean *T_leaf_* during the 2024-2025 austral summer, relative to control trees (F_1,9_ = 6.29, *p* = 0.033; Fig. 6a), with an average difference of 1.4 °C. Leaf critical temperature (*T_crit_*) was equivalent between irrigated and control trees (48.2 vs. 47.8 °C, respectively; F_1,6_ = 0.27, *p* = 0.61). The thermal safety margin of irrigated trees was an average of 1.7 °C higher than control trees (F_1,9_ = 6.2, *p* = 0.034) on a hot day (28 January 2025, maximum *T_air_* = 40.2 °C) in mid-summer. Leaf photosynthetic efficiency significantly declined with increasing *T_leaf_* (R^2^ = 0.45, *p* < 0.001; Fig. 6b), although the difference between treatments was not significant (F_1,10_ = 1.14, *p* = 0.31). *T_air_* was significantly correlated with *T_leaf_* (R^2^ = 0.71, *p* < 0.001; Fig. 6c), but the relationship was not different between treatments. Leaf temperature exceeded air temperature on the surveyed dates by 2.4-10.1 °C but was more similar to *T_air_* on extreme days approaching 40 °C, especially for irrigated trees (Fig. 6c). On an extremely hot and dry day in early autumn, on 13 March 2025 (maximum *T_air_* = 41.1 °C, soil VWC = 7.3%), our thermal infrared images confirmed that the crown of control trees (Fig. S6f) reached a maximum crown temperature of 43.2-47.7 °C, while irrigated trees peaked at 39.9-41.8 °C (Fig. S6d).

**Figure 6.**
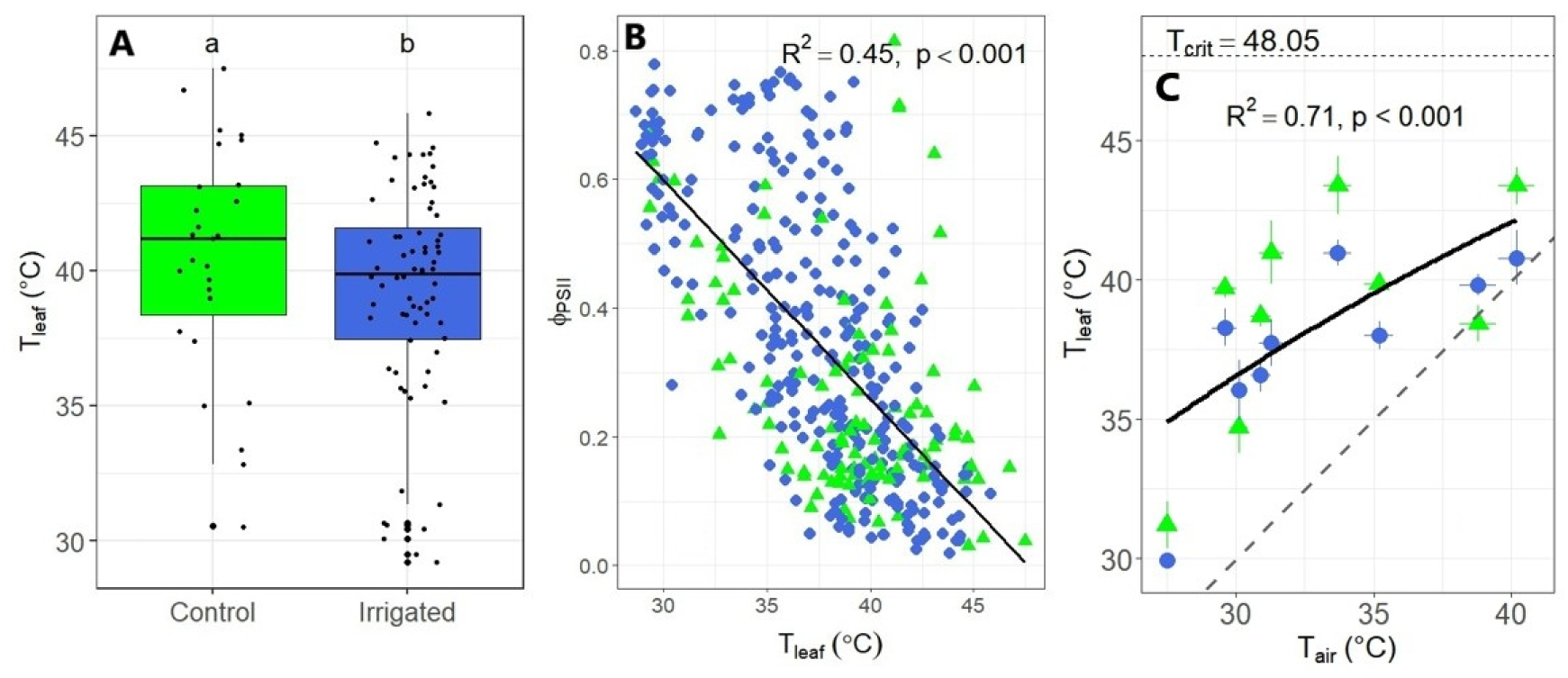
(A) Comparison of peak leaf temperature (*T_leaf_*) between irrigated (N = 8 trees) and control trees (N = 3 trees) of *Lophostemon confertus* in the western Sydney suburb of Tregear, NSW, Australia over the austral 2024-2025 summer on averaged (N = 4) leaves per tree. Error bars represent ± standard error of the mean; whiskers indicate 1.5 × interquartile range. Smaller points indicate raw data, while bigger points the minimum values. (B) Linear relationship between leaf photosynthetic efficiency of photosystem II (Φ_PSII_) and leaf temperature (*T_leaf_*). (C) Nonlinear relationship between air temperature (*T_air_*), as recorded by the weather station on site during the measurement time (13:00 to 15:00), and *T_leaf_*. The horizontal dashed line denotes the species mean leaf critical temperature. Black line is *T_leaf_*=nls(*T_air_*). Grey dashed line is the 1:1 line. Irrigated trees are coloured blue and represented in circles, controls in green and triangles. Error bars represent SE. Means not connected by the same letter are significantly different (Tukey honestly significant difference, *p* < 0.05).

## 4 Discussion

Cities can impose harsh conditions for tree establishment and growth due to the combined effects of elevated urban temperatures, high irradiance, and reduced access to water, which intensify cumulative thermal stress on vegetation during summer heatwaves (Oke, 1991; Gillner et al., 2015; Rahman et al., 2017; Marchin et al., 2022a, 2025). Under such conditions, adequate water availability is critical to sustain transpiration and enable evaporative leaf cooling in urban trees (Winbourne et al., 2020; Tams et al., 2024), although the capacity of different species to sustain transpiration and mitigate thermal stress varies with species-specific water use strategies and microclimatic conditions (Breen et al., 2004; Moore, 2013). Our study evaluated the benefits of a passive irrigation system that collected and stored rainwater for young, establishing broadleaf evergreen trees (*L. confertus*) along a residential street in western Sydney, Australia. Brush box is native to Sydney’s climate and was relatively tolerant (i.e., crown dieback in most trees <25%) to the transient periods of moderate heat and drought stress experienced during our study. This commonly planted species can tolerate extreme heatwaves when irrigated (Esperon-Rodriguez et al. 2020), but its tolerance may decline under water limitation, with *L. confertus* classified as heat-intolerant under low-rainfall conditions despite relatively modest mean dieback (Marchin et al. 2022). At the peak of summer, the soil REW at 45-cm depth was 13% higher near irrigated trees than by the control trees (Fig. S3b; Table S1). The greater water access allowed irrigated trees to maintain greater leaf water potentials (*Ψ_pre_* and *Ψ_mid_*) than control trees (Fig. 3, 4), indicating lower hydraulic stress that likely contributed to their lower crown dieback (Fig. 1). Irrigated trees also exhibited greater *g_s_* on dry summer days (Fig. 5b), along with lower leaf temperatures (Fig. 6), despite having larger leaves (Fig. 2), which ultimately resulted in greater latent heat flux and a cooler canopy. By maintaining transpiration on hot, dry days, irrigated trees effectively mitigated thermal stress as evidenced by their greater crown survival. Irrigated trees had larger canopy area in their second year after planting but overall, their growth rates were similar to control trees. These results agree with findings from other PI systems, where additional water supply had little effect on the growth of moderate to drought-tolerant species (Luketich, Papuga & Crimmins 2018). Instead, passive irrigation helped mitigate heat and drought stress in urban trees, thereby increasing their cooling capacity and strengthening the climate resilience of western Sydney.

### 4.1 Passive irrigation enhanced crown survival without increasing growth

Passive irrigation systems have been implemented in Adelaide, Sydney, and Melbourne in Australia (EPA 2013; Szota et al., 2019; Gleeson et al., 2022) and other cities worldwide (Bartens et al., 2009; Morgenroth & Buchan 2009; Ow & Chan, 2021; Dowtin et al., 2023). Passive irrigation has the potential to both benefit urban trees and, when implemented at scale, to mitigate the risk of flooding by capturing stormwater (Szota et al., 2019; Thom et al., 2022). Street tree growth and resilience may be enhanced by increasing access to water, while moving stormwater into storage pits allows for increased soil water infiltration on city streets (Bartens et al., 2008) and reduced flow into stormwater systems (Smith et al., 2024). Our study species, *L. confertus* (brush box), has been suggested to perform well in passive irrigation systems, as it has high transpiration and moderate recovery of transpiration following drought (Thom et al. 2022, 2022a). The effects of passive irrigation on canopy physiological function and health in our research were clear and most pronounced during hot, dry days (Fig. 5b). Greater water access throughout the summer supported leaf thermoregulation to consistently mitigate cumulative heat stress due to repeated hot days, which increased crown survival in a hot, dry summer in western Sydney. Our results suggest that urban trees can be vulnerable to transient heat events (*T_air_* > 40 °C) and that even well-adapted native species benefit from passive irrigation during peak stress periods.

Passive irrigation (PI) systems have shown variable effects on urban tree growth, with responses strongly dependent on the growth strategy of individual species, background climate context, and the timing and length of observations (Morgenroth & Buchan 2009; Luketich, Papuga & Crimmins 2018). In Singapore, passively irrigated mature *Calophyllum inophyllum* (Alexandrian laurel) trees had 3 to 6 times greater growth than controls (Ow & Chan, 2021). In Adelaide, passively irrigated *Melia azedarach* (Chinaberry tree) saplings showed 60-65% greater height growth and trunk expansion (DBH), compared to non-irrigated trees (Gleeson et al., 2022). Growth rates were 44% higher in irrigated *Acer campestre* (field maple) saplings in Melbourne during the first two establishment years (Thom et al., 2022). In established trees, however, PI systems produced negligible growth responses across multiple species in Melbourne (Szota et al. 2019). Contrary to our hypothesis, our study did not show any sustained difference in growth between treatments during the early establishment years (Table 2), although there was 37% greater canopy expansion of irrigated trees during a year with below-average rainfall (MAP_2022-2023_ = 740mm compared to MAP_1975-2025_ = 930mm). Overall, growth benefits of PI systems may be limited in some species, perhaps due to slower growth rates (Ren et al., 2023). Additional variability may arise from differences in climate context, with stronger effects expected in chronically water-limited cities or extreme drought years, as well as from system design and how it influences soil moisture redistribution and root-zone hydrology (Scharenbroch et al., 2016; Gleeson et al., 2022; Thom et al., 2022). Future studies should focus on identifying which species are best suited for passive irrigation systems (Thom et al. 2022a), and testing their performance on urban streets, to maximize tree performance and ecosystem services in hot suburbs.

### 4.2 Irrigated trees had improved leaf water status and thermoregulation

Irrigated trees benefitted from enhanced water availability, with mean daily REW being on average 13% greater at 45-cm depth, maintaining less negative *Ψ_pre_* and *Ψ_mid_* compared to controls (Fig. S3b; Table S1). While *Ψ_pre_* is more suited to track tree water availability (Marchin et al., 2025), differences in *Ψ_mid_* suggest that irrigated trees operated under reduced hydraulic stress during peak atmospheric demand (Fig. 4b). When interpreted through a hydraulic safety margin framework, control trees more frequently exceeded their turgor loss point, or wilting point. Repeated or prolonged excursions beyond TLP are associated with a greater risk of hydraulic failure resulting in desiccation damage and canopy dieback (Bartlett et al., 2016b). These dynamics are likely exacerbated in urban environments, where evaporative demand and hydraulic strain are intensified by impervious surfaces and urban heat (Nowak, 2010). During an extreme summer in western Sydney, *L. confertus* was vulnerable to hotter drought with a mean relative dieback rate of 32 ± 8% (Marchin et al., 2025). Our findings suggest that passive irrigation can effectively mitigate drought stress in this species, as irrigated trees exhibited improved hydraulic function, and reduced crown dieback (Fig. 1).

Leaf temperature depends not only on transpiration rate but also on morphological traits such as leaf size and canopy structure (Vogel et al., 2009; Leuzinger et al., 2010, Leigh et al., 2012, 2017; Urban et al., 2017). Leaf temperatures under heatwave conditions can exceed critical thresholds when water availability is limited (Lambers et al., 2008). Elevated leaf temperatures can occur due to limited transpiration under conditions of reduced hydration during water deficit (Taiz et al., 1999). In agreement with our hypothesis, irrigated trees had lower *T_leaf_* than controls (Fig. 6a). Across all study trees, *T_leaf_* was more closely coupled with *T_air_* on an extremely hot day (*T_air_*>40 °C) than on average summer days (Fig. 6c). As a result, irrigated trees exhibited greater transpiration rates on hot and dry days and experienced greater crown cooling, relative to controls (Fig. 3, S5). On three hot and dry days, irrigated trees had higher mean *g_s_* and greater thermal safety margins than controls (Fig. 5b). Increased *g_s_* functions to cool leaves during heatwaves, enabling trees to mitigate the effects of extreme heat (Bachofen et al., 2025). On days after recent rainfall, when more water was available for all trees, irrigated trees instead prioritised stomatal closure, resulting in similar rates of *g_s_* to control trees (Fig. 5c). Three irrigated trees exhibited consistently higher *g_s_* rates than the other irrigated trees across the 2024-2025 austral summer (Fig. S4c), although their *Ψ_pre_* did not differ from the other trees. Similarly, irrigated street trees with similar *Ψ_mid_* had higher *g_s_* than nearby park trees in autumn (Fig. 3). One potential explanation is microclimatic differences in urban temperatures among our study trees – one limitation of our study was an inability to control the spatial location of control, irrigated, and park trees. Regardless, our results identify that irrigated trees were able to maintain higher *Ψ_mid_* and higher *g_s_* than unirrigated controls and/or park trees at certain times of the year (Fig. 3).

Thermal imaging offers a rapid and easily repeatable monitoring of tree canopy temperature, which can be used to indicate plant water status (Carrasco-Benavides et al., 2020; Asawa et al., 2024). This non-invasive tool could be used to apply targeted watering to avoid heat and water stress in trees that do not receive regular irrigation (Asawa et al., 2024; Javadian et al., 2024; Wilkening and Feng, 2025). It could also be used to assess degree of canopy cooling and identify heat-resilient species (Jones, 1999; Leuzinger and Körner, 2007; Moser et al., 2017; Munné-Bosch et al., 2021; Pfautsch et al., 2021). Notably, even low-resolution thermal cameras have been shown to provide reliable water-stress assessments comparable to higher-end systems (Fig. S6; Carrasco-Benavides et al., 2020).

## 5 Conclusions

Rising temperatures, particularly during summer heatwaves, pose substantial risk of thermal stress to urban trees. Adequate water supply remains critical for urban trees to avoid heat and drought stress. Our study has demonstrated that passive irrigation systems can improve tree water access and increase leaf water potential in the drought-tolerant native Australian species, *Lophostemon confertus*, allowing continued transpiration and supporting greater canopy survival. Although differences in *g_s_* were restricted to the hottest and driest days, irrigated trees maintained lower leaf temperatures and enhanced physiological performance during critical periods of thermal stress. These findings suggest that passive irrigation can enhance the resilience of urban trees by buffering against episodic heat stress, although growth may not be promoted in tolerant species. In increasingly hot urban environments, future research should focus on a greater understanding of which tree species, locations and climate conditions are best suited for the installation of PI systems to maximise cost effectiveness and provision of ecosystem services, including cooling to mitigate urban heat.

## 6 Data Availability

The raw data supporting the conclusions of this article are publicly available at Figshare, https://doi.org/10.6084/m9.figshare.32112781.

## 7 Acknowledgements

The authors thank Blacktown City Council for building and maintaining the study site, and Chris Jones for his assistance; the views expressed herein are those of the authors and are not necessarily those of Blacktown City Council.

## Author Contributions

Davide Siclari: Investigation, Formal analysis, Visualization, Writing – original draft. Paul D. Rymer: Conceptualization, Writing – review & editing, Funding acquisition. Mark G. Tjoelker: Writing – review & editing. Sebastian Pfautsch: Writing – review & editing.

Chathurika Perera: Methodology, Writing – review & editing. Renée M. Marchin: Conceptualization, Writing – review & editing, Funding acquisition.

