## Supplementary Data for "Passive irrigation increases transpiration and reduces heat stress for urban *Lophostemon confertus* trees in metropolitan Sydney"

Table S1. Maximum daily air temperature (*T_air_* °C), vapour pressure deficit (VPD, kPa), maximum daily wind speed (km/h), treatment-specific relative extractable water content (REW, based on measured soil VWC at 45 cm depth), and estimated soil moisture conditions (Australian Landscape Water Balance Model), on the four measurement days during austral autumn 2024 (shaded grey) and the nine measurement days during the austral 2024-2025 summer. Data source: weather station on site.

| **Date** | **T_air_ (°C)** | **VPD (kPa)** | **Wind speed (km/h)** | **REW Irrigated** | **REW Controls** | **Estimated Soil Moisture** |
| --- | --- | --- | --- | --- | --- | --- |
| 2024-03-14 | 31.4 | 1.84 | 15.32 | - | - | Very Dry |
| 2024-04-12 | 22.5 | 1.01 | 9.12 | - | - | Saturated |
| 2024-05-14 | 19.1 | 1.06 | 11.21 | - | - | Average/Moist |
| 2024-06-18 | 20.4 | 1.25 | 7.09 | - | - | Moist |
| 2024-11-27 | 38.8 | 6.90 | 18.79 | 0.33 | 0.34 | - |
| 2024-12-05 | 27.5 | 3.65 | 8.93 | 0.77 | 0.53 | - |
| 2024-12-27 | 35.2 | 5.67 | 18.14 | 0.22 | 0.09 | - |
| 2025-01-05 | 33.7 | 5.21 | 10.66 | 0.04 | 0.01 | - |
| 2025-01-13 | 30.1 | 4.24 | 9.58 | 0.96 | 0.95 | - |
| 2025-01-22 | 31.3 | 4.54 | 9.65 | NA | NA | - |
| 2025-01-28 | 40.2 | 7.43 | 19.98 | NA | NA | - |
| 2025-02-05 | 34.3 | 5.38 | 13.90 | NA | NA | - |
| 2025-02-07 | 29.9 | 4.19 | 11.05 | NA | NA | - |

Table S2. Growth parameters (expressed in cm), including height, canopy width and length, diameter at breast height (DBH, 1.3m) and basal diameter one year after establishment (23 February 2022) between irrigated and control (shaded grey) trees of *Lophostemon confertus* in Wilkes Crescent, Tregear, Blacktown NSW, AU.

| **Tree ID** | **Height** | **Canopy Width** | **Canopy Length** | **DBH** | **Basal** |
| --- | --- | --- | --- | --- | --- |
| 1 | 309 | 162 | 156 | 3.12 | 6.4 |
| 2 | 386 | 200 | 190 | 4.5 | 6.1 |
| 3 | 294 | 149 | 150 | 3.25 | 4.8 |
| 4 | 312 | 145 | 152 | 3.45 | 5.1 |
| 5 | 284 | 151 | 150 | 2.65 | 4.1 |
| 6 | 325 | 162 | 162 | 3.75 | 5.6 |
| 7 | 324 | 162 | 159 | 3.2 | 5 |
| 8 | 392 | 210 | 225 | 4.15 | 6 |
| 9 | 313 | 133 | 136 | 3.15 | 6 |
| 10 | 301 | 172 | 181 | 3 | 5.8 |
| 11 | 305 | 135 | 150 | 3.45 | 5.1 |
| 12 | 401 | 180 | 185 | 3.65 | 5.5 |
| 13 | 328 | 192 | 185 | 3.55 | 5 |
| 14 | 339 | 195 | 184 | 4.25 | 6.3 |
| 15 | 286 | 155 | 149 | 2.9 | 4.8 |
| 16 | 381 | 169 | 181 | 4.35 | 6.6 |
| 17 | 262 | 157 | 168 | 2.5 | 4.4 |
| 18 | 363 | 168 | 166 | 3 | 5.5 |
| 19 | 365 | 200 | 215 | 3.6 | 7 |
| 20 | 333 | 170 | 184 | 3.4 | 5.9 |
| 21 | 261 | 176 | 147 | 2.6 | 5.4 |
| 22 | 385 | 204 | 195 | 4.75 | 7 |
| 23 | 278 | 204 | 213 | 4 | 7 |
| 24 | 323 | 175 | 170 | 3.45 | 6 |
| 25 | 315 | 135 | 138 | 3.35 | 4.4 |

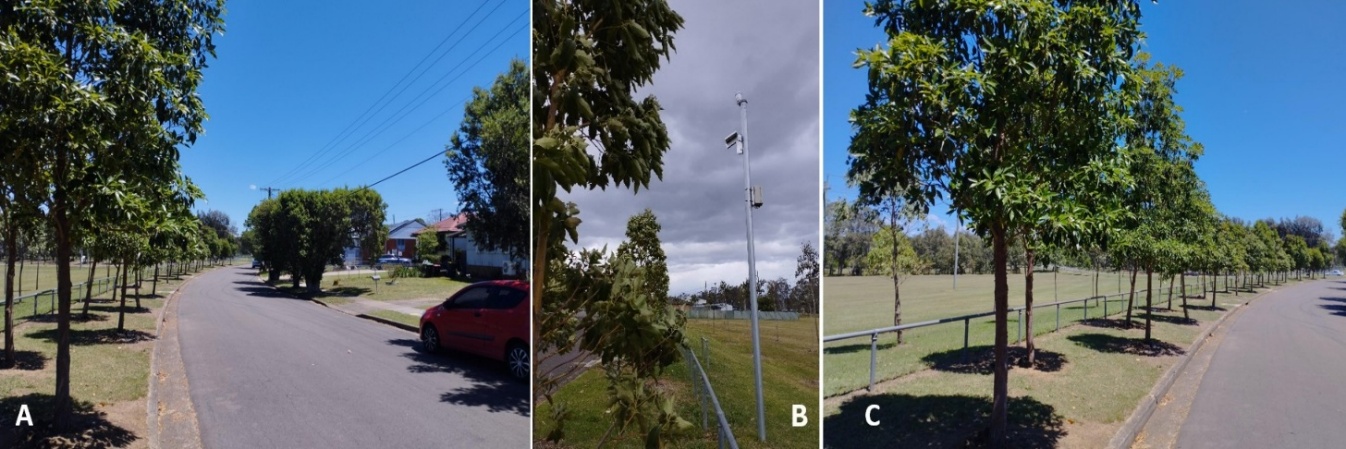

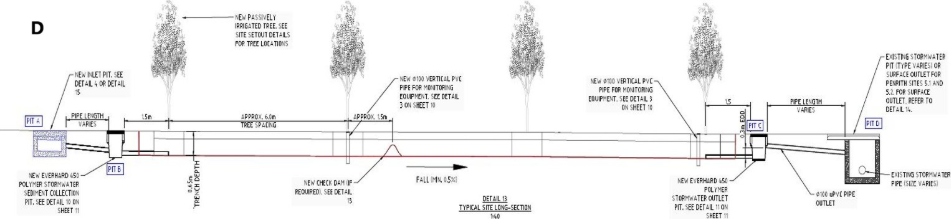

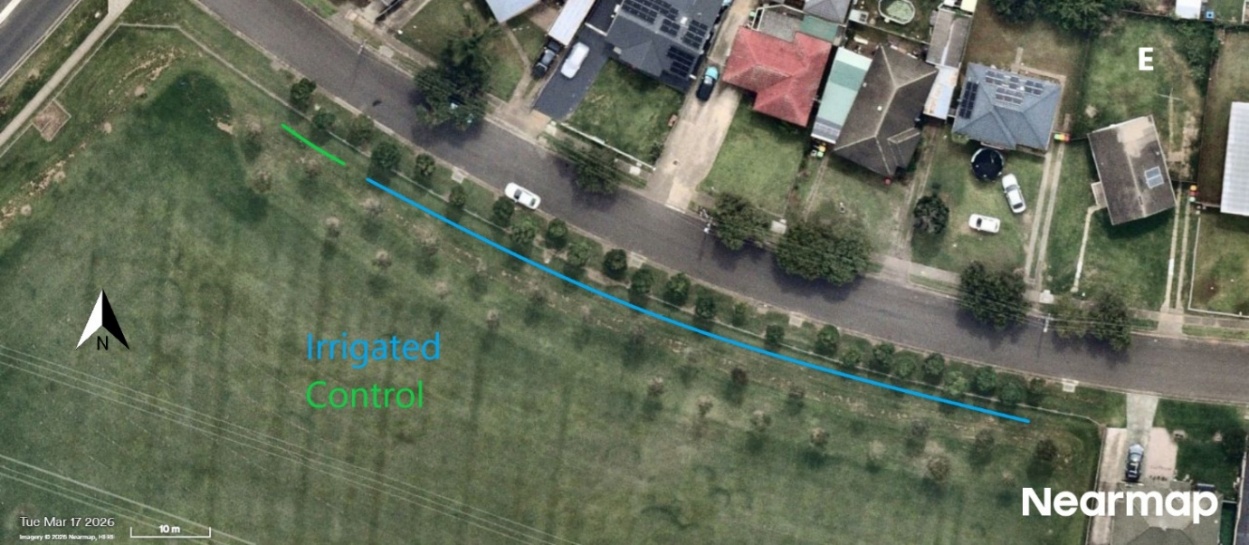

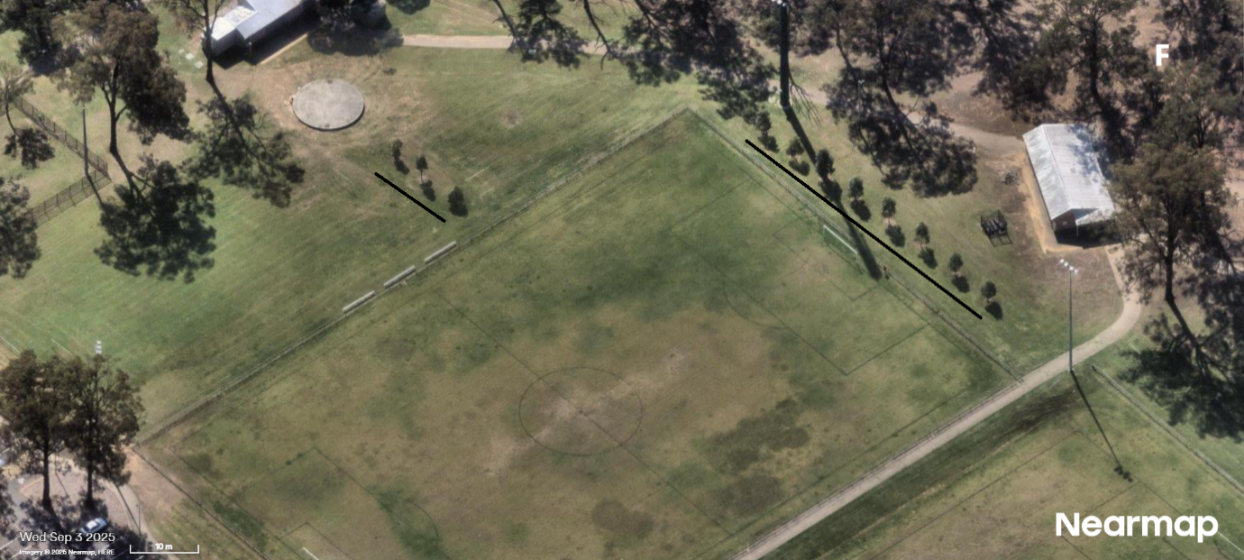
Figure S1. (A) The study site located at Wilkes Crescent, Tregear, Blacktown, NSW, Australia, with view of the residential houses on the right. (B) Weather station installed on site. (C) The full line of 25 *Lophostemon confertus* trees with the recreation area on the left. (D) A schematic drawing of the passive irrigation system with 4 trees. E) Map of the site with irrigated trees highlighted in blue and control plants in green. (F) The north-oriented map of the park control trees planted in Shanes Park Reserve, Blacktown (4 km away from the study site).

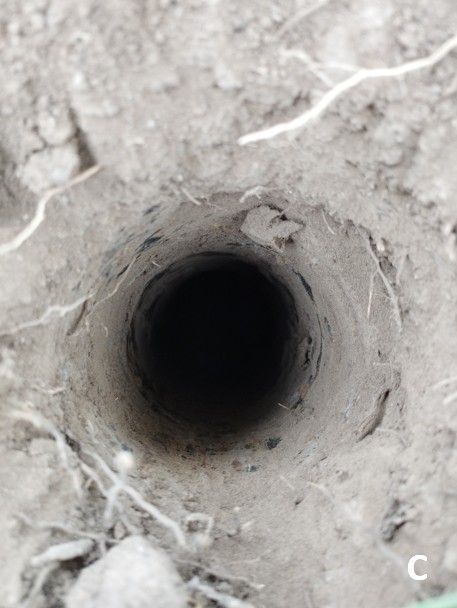

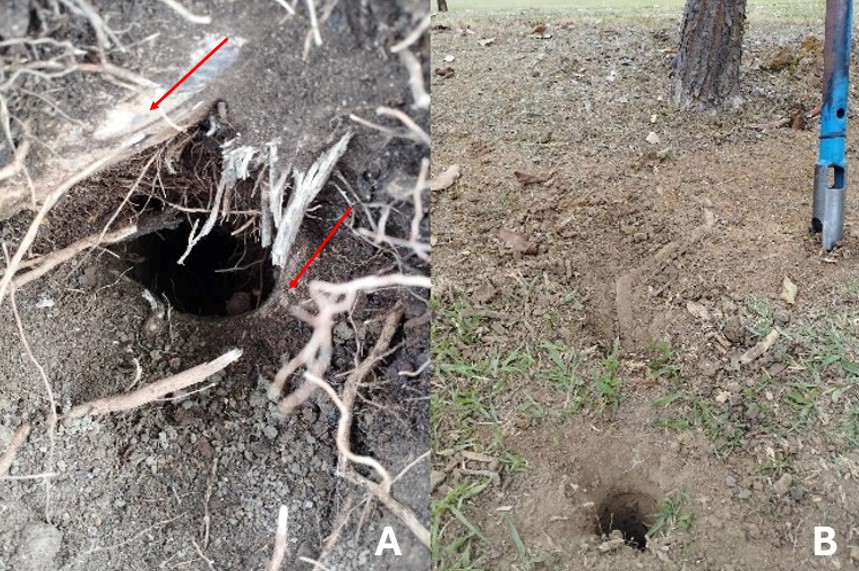
Figure S2. (A) An initial soil core near the base of an irrigated *Lophostemon confertus* tree revealed two main lateral roots, highlighted by two red arrows. Roots (diam.=3-4cm) were traced and excavated along their trajectory through the surface soil. (B) A second exploratory soil core was dug 15-20 cm further along the assumed root path. (C) If no root continuation was observed at a depth of 50 cm, it was inferred that the root descended vertically into the passive irrigation (PI) trench, located at a depth of 0.65 m. Investigative cores were conducted on three experimental trees with identical results.

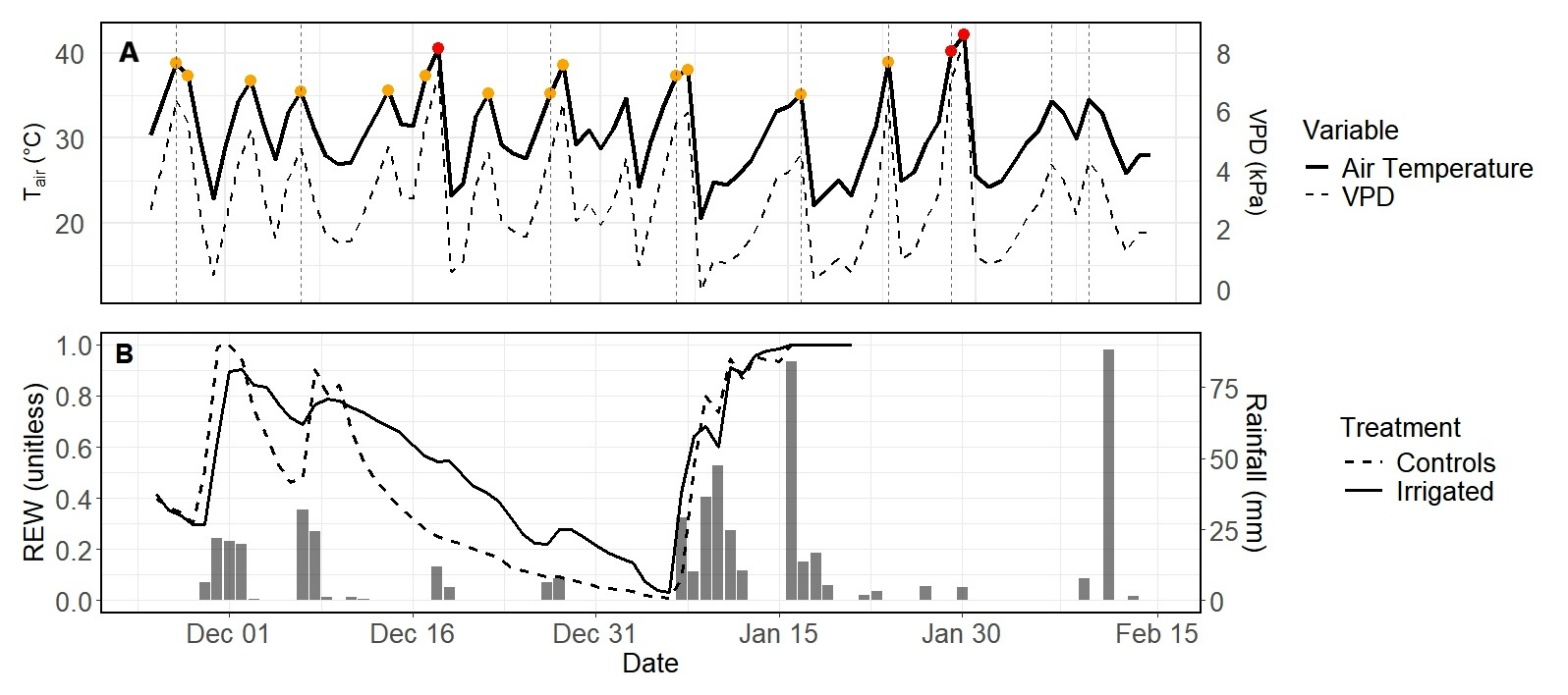
Figure S3. Summer climatic conditions, soil water availability and rainfall during the austral 2024–2025 summer in the western Sydney suburb of Tregear, NSW, Australia. (A) Daily maximum air temperature (solid line) and vapour pressure deficit (VPD; dashed line) recorded on site in Tregear during the austral 2024-2025 summer. Vertical dashed lines indicate measurement dates, orange points are hot days (>35 °C), and red points are extreme days (>40 °C). (B) Mean relative extractable water content (REW, unitless) at a soil depth of 45 cm is compared shown for between irrigated (solid line, N = 3 soil moisture sensors) and control trees (dashed line, N = 2 soil moisture sensors). Daily rainfall is also shown (grey bars).

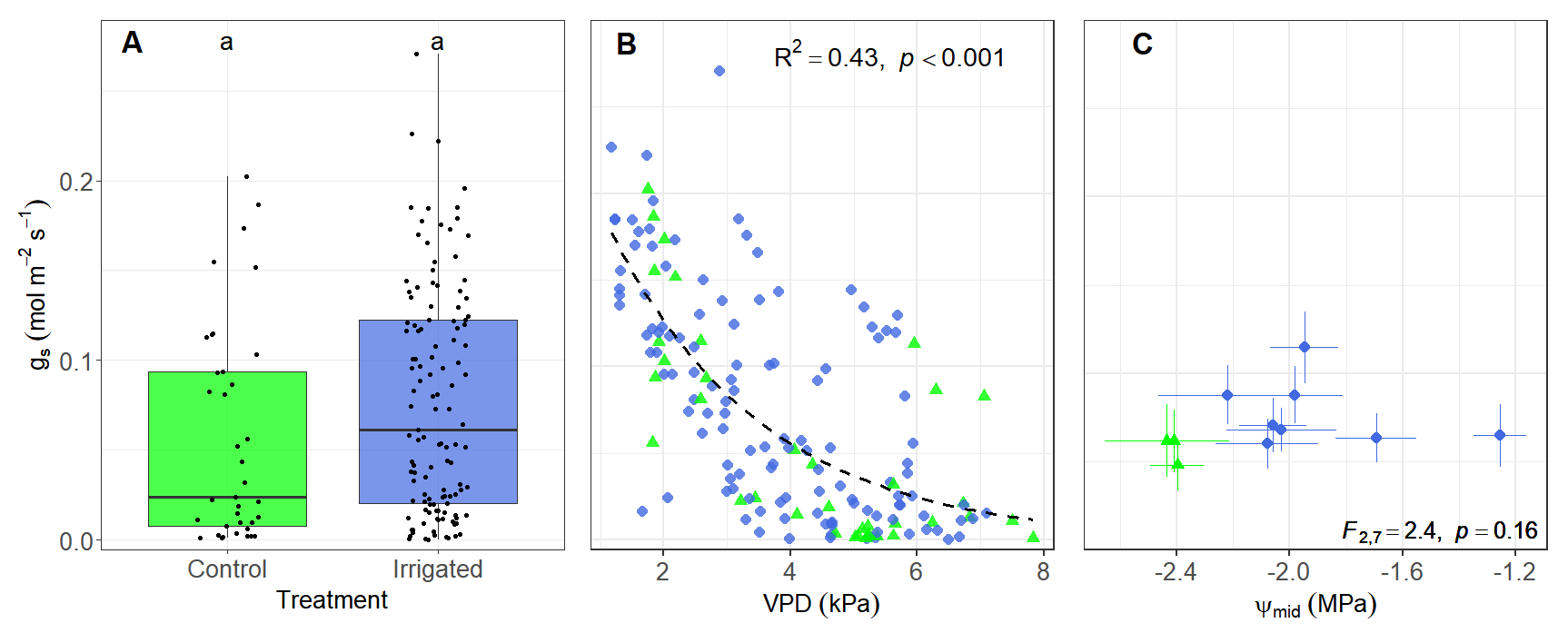
Figure S4. (A) Comparison of *g_s_* between irrigated (N = 8 trees) and control trees (N = 3 trees) of *Lophostemon confertus* in the western Sydney suburb of Blacktown, NSW, Australia. Each point is the mean of 4 leaves per tree per sampling date in the austral 2024-2025 summer. Error bars represent ± standard error of the mean; whiskers indicate 1.5 × interquartile range. (B) Daily mean stomatal conductance and vapour pressure deficit (VPD), with the fitted non-linear relationship (dashed line) *g_s_*=ln(VPD). (C) Mean tree-level *g_s_* and midday leaf water potential (*Ψ_mid_*). Irrigated trees are coloured blue and represented in circles, controls in green and triangles. Error bars represent SE. Means not connected by the same letter are significantly different (Tukey honestly significant difference, p<0.05).

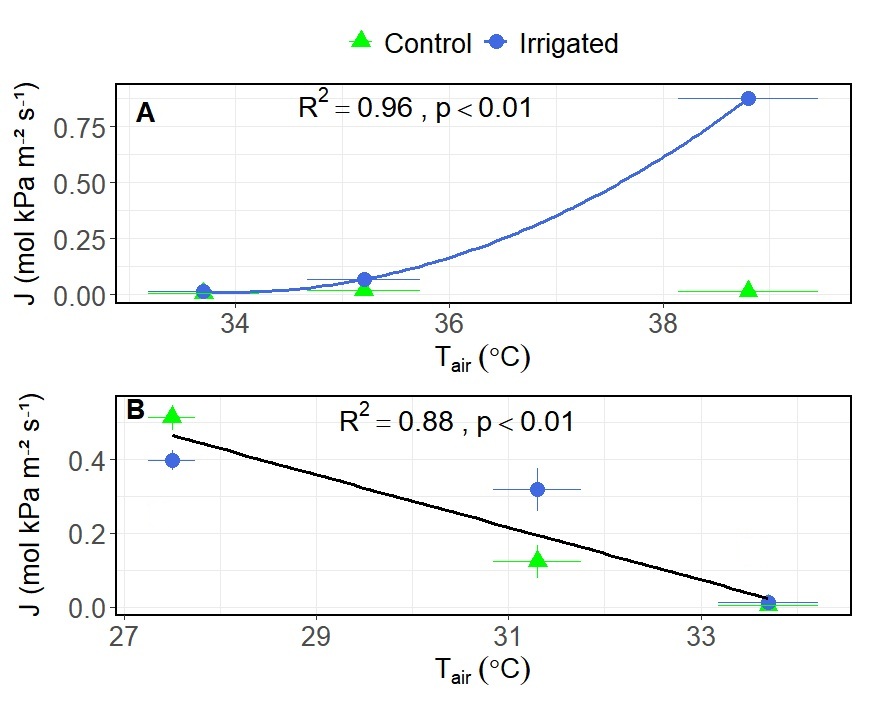

Figure S5. Relationship between transpiration (*J=g_s_ x VPD*) and air temperature (*T_air_*) calculated on irrigated (N = 8) and control trees (N = 3) of *Lophostemon confertus* in the western Sydney suburb of Blacktown, NSW, Australia on (A) hotter, dry days (VPD > 5 kPa; 5 December 2024, 13, 22 January 2025) *T*=ln(*T_air_*) versus (B) wetter days (VPD < 3 kPa; 27 November , 27 December 2024, 5 January 2025) during the austral 2024-2025 summer. Irrigated trees are coloured blue and represented in circles, controls in green and triangles. Error bars represent SE.

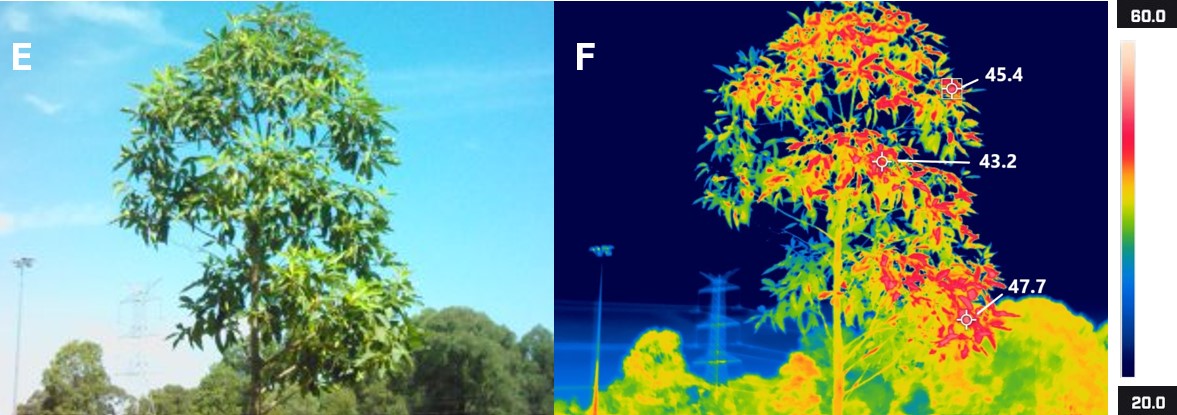

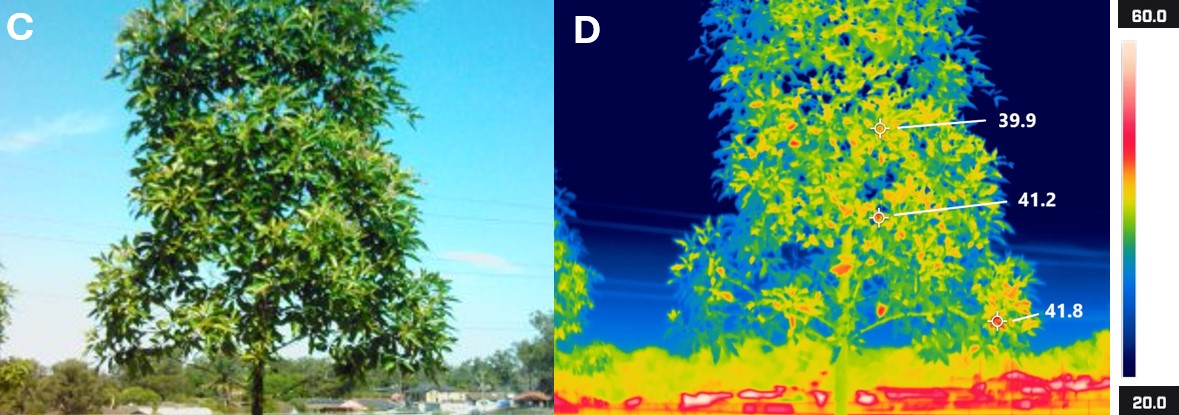

Figure S6. Comparison between RGB (left) and thermal infrared (right) images. (A), (B) The entire line of *Lophostemon confertus* trees along the passive irrigation trench, highlighting surface temperatures across various sunlit and shaded materials, scaled from 20 to 80 °C. (C), (D) An irrigated *L. confertus* tree and (E), (F) a control *L. confertus* tree without access to passive irrigation. All images were captured on a hot and dry day in early autumn, 13 March 2025 (maximum air temperature = 41.1 °C; VWC=7.3 %), on the north-facing side of the street, fully exposed to sunlight. Tree orientation was confirmed by the consistent appearance of overhead powerlines in the background. In the thermal images, hotter areas appear in red tones and cooler areas in blue. White crosshair markers denote selected pixels in the thermal images; temperature values are in °C. C, D, E and F are scaled 20 to 60 °C.
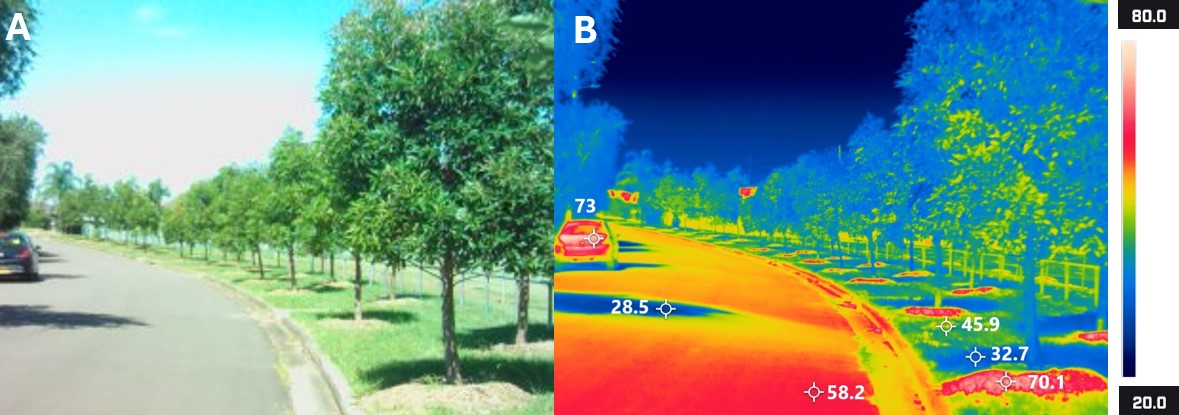
